# Micro-hotspots of Risk in Urban Cholera Epidemics

**DOI:** 10.1101/248476

**Authors:** Andrew S. Azman, Francisco J. Luquero, Henrik Salje, Nathan Naibei Mbaïbardoum, Ngandwe Adalbert, Mohammad Ali, Enrico Bertuzzo, Flavio Finger, Brahima Toure, Louis Albert Massing, Romain Ramazani, Amélie Cardon, Bansaga Saga, Maya Allan, David Olson, Jerome Leglise, Klaudia Porten, Justin Lessler

## Abstract

Targeted interventions have been delivered to neighbors of cholera cases in epidemic responses in Haiti and Africa despite little evidence supporting impact. Using data from urban epidemics in Chad and D.R. Congo we estimate the size and extent of spatiotemporal zones of increased cholera risk around cases. In both cities, we found zones of increased risk of at least 200-meters during the 5-days immediately following case presentation to a clinic. Risk was highest for those living closest to cases and diminished in time and space similarly across settings. These results provide a rational basis for targeting interventions, if delivered rapidly.

## Background

Cholera epidemics in sub-Saharan Africa produce a large proportion of global cholera-mortality and continue to wreak havoc on already fragile nations.^1-3^ Targeting cholera interventions to transmission hotspots, or areas of elevated transmission intensity in urban areas may be the best control strategy when resources are constrained.^4, 5^ In recent years, rapid response teams have been proposed as an important way to fight cholera in cholera prone countries. These teams can quickly provide emergency water, sanitation and hygiene interventions (e.g., point of use water treatment and basic hygiene educational materials), and sometimes oral cholera vaccine to neighbors of cholera cases.^6, 7^ However, limited evidence exists regarding the impact or optimal spatial scale of these targeted interventions.

Cholera transmission is thought to occur through two modes of exposure: (1) environmentally-mediated exposure, often because fecal contamination in the broader environment (e.g., water sources), and (2) direct exposure, through more direct contact with bacteria (e.g., contaminated food).^8^ The mix of environmentally-mediated and direct exposure shapes the spatiotemporal distribution of cases within an epidemic. Evidence from Bangladesh and other locations have shown that direct transmission plays an important role in cholera transmission leading to elevated risk when residing close to an incident case, and that cholera risk factors can cluster at small spatial scales.^9, 10^ Due to limited high-resolution spatially-explicit surveillance data in epidemics, especially in sub-Saharan Africa, little is known about the mix of exposure processes which shape the size and duration of zones of increased risk around incident cases.

Characterizing the small-scale spatiotemporal distribution of cholera cases in epidemics can provide new and useful insight into the mechanisms of transmission ultimately highlighting a path for efficient targeted cholera control. Here we use high-resolution data from epidemics in two African cities, thousands of kilometers apart, to illustrate spatiotemporal windows of increased cholera risk and discuss their implications for targeting interventions at neighbors of incident cholera cases.

## Methods

### Study Setting

Chad has experienced cholera outbreaks at least once every 4-years since the 1990’s. Médecins Sans Frontières (MSF) assisted the Chad Ministry of Health to respond to a cholera outbreak that started in mid-April 2011 in which most cases occurred in the capital, N’djamena, home to nearly one million people. On 22-June-2011 the MSF team, with the assistance of other collaborating agencies, began systematically collecting household GPS coordinates through a home visit to each suspected cholera case presenting at one of the official cholera treatment centers/units in N’Djamena. As the number of cases per day began to rapidly increase in early October, MSF modified their protocol to collect household coordinates for one out of every three cases.

Kalemie is located on Lake Tanganyika in eastern Democratic Republic of Congo and serves as a large urban trading center for the region. Cholera tends to occur annually in Kalemie with a seasonal peak within the last few months of the year. In Kalemie, MSF has worked with the Ministry of Health on comprehensive cholera prevention and control strategies since 2008. From Jan-2012 to Jan-2013, MSF and the MoH collected the household coordinates for each suspected cholera case seeking care at the main diarrhea treatment center in Kalemie, Centre de Traitement de Maladie Diarrhéique.

In both settings, suspected cases were defined using a modified WHO case definition (acute watery diarrhea regardless of age). Teams in both countries were trained in the use of the GPS devices to ensure stable and accurate readings.

### Statistical Approach

To characterize the spatiotemporal clustering of cases, we calculated the *τ*-statistic, a global clustering statistic estimating the relative risk of the next case occurring at a distance *d*, within *t* days after a suspected case presents at a health facility compared to the risk of the next case occurring anywhere in the population during the same period. We calculated the *τ*-statistic using the `IDSpatialStats` package^11^, with a 50-meter moving window estimated every 10-meters. This statistic is robust to heterogeneities in the spatial structure of the underlying population and cases missing at random.^11, 12^ We calculated 95% confidence intervals as the 2.5^th^ and 97.5^th^ quantiles from 1,000 bootstrap replicates.

We focused on estimates of this relative risk (*τ*) at distances up to 500-meters from a primary case and within 5-day windows up to 30-days after a ‘primary’ case presented to a facility. We used a small spatial moving window so that our estimates of *τ* for clustering at one distance to have minimal effect on other distances, therefore our estimates are not necessarily smooth, nor monotonic. We considered the zones of increased risk around incident cases to extend until the 95% confidence intervals cross unity for at least two consecutive points (i.e., 20 consecutive meters). Since this classification may underestimate the extent of the zones of increased risk due to small sample size, we calculated the median (and 2.5^th^ and 97.5^th^ quantiles) distance at which *τ* dropped below 1.2 (e.g., a minimum 20% elevated risk) for each bootstrap as an alternative measure. Since it is unlikely that a targeted public health response can be mounted the same day a case seeks care (day 0), we also estimated the zones of increased risk around incident cases excluding the day of case presentation.

## Results

In Kalemie, DRC, household coordinates were successfully recorded for all 1,146 suspected cholera cases reporting to the main diarrhea treatment center from January 2013 through January 2014. In N’djamena, household coordinates were recorded for 1,692 of the 4,359 suspected cases reporting to healthcare facilities within the city. All case households were visited before August-2011 and one in three randomly selected case-households visited from August through the end of the outbreak in December due to logistical constraints.

### The first five days

Within the first five days after a suspected cholera case presented for care, we found that the zone of increased cholera risk extended to at least 210m from the home of the suspected cases in Kalemie and 330m in N’Djamena. Zones of increased risk estimated using a bootstrap approach were similar (Table S1). Those living within 20m of another case (including those living in the same household) had a 44.5-fold (Kalemie; 95%CI 30.1-68.3) higher risk than the general population of becoming a cholera cases within five days of the index case in Kalemie, and a 32.4-fold (95%CI 25.3-41.0) higher risk in N’Djamena (Figures 1A-B). Those living 75-125m from a case had 1.9 (Kalemie; 95%CI 1.1-3.0) and 3.9 (N’Djamena; 95%CI 2.7-5.4) times the cholera risk in the five days after the initial case compared to those elsewhere in the city.

**Figure 1.**
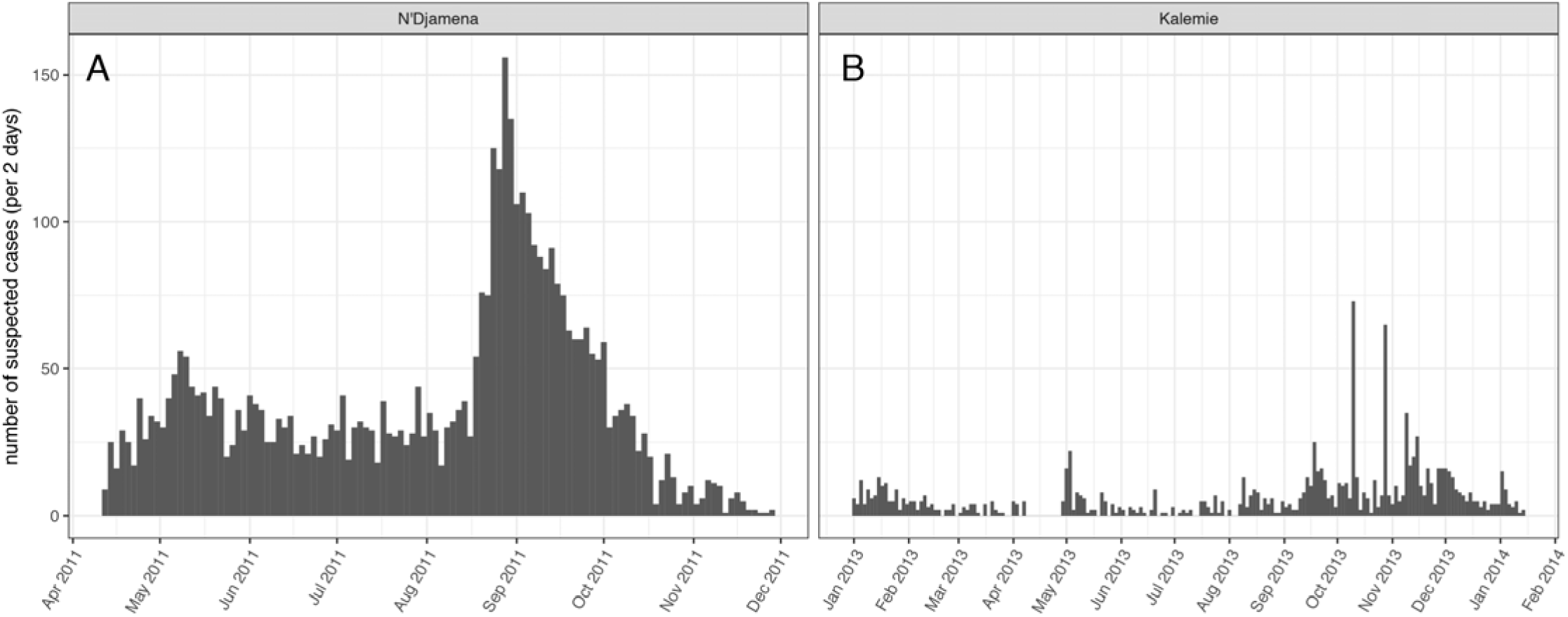
Epidemic curves from Ndjamena, Chad and Kalemie, Democratic Republic of Congo (A and B).

**Figure 2.**
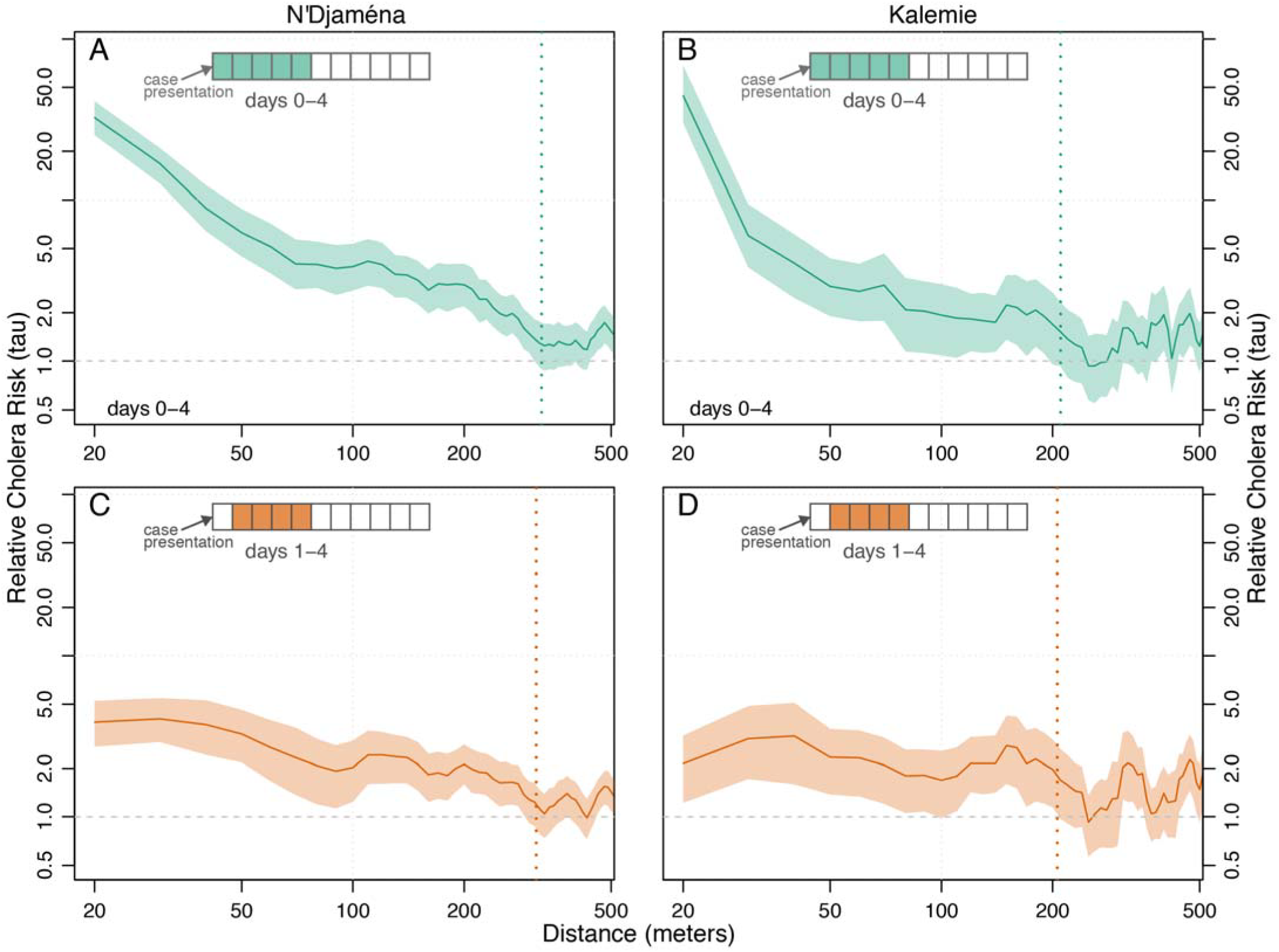
Estimates of the relative risk of the next cholera case being within a specific distance to another case (x-axis) within either days 0-4 (green, A-B) or days 1-4 (orange, C-D) compared to the risk of the case occurring anywhere in the population. Dashed lines represent the spatial extent of the zones of increased risk as defined by the first point at which the 95% confidence intervals cross unity over a 20-meter interval (i.e., over two consecutive 10-meter points).

Within these first five days after a case presented for care, the most elevated risk occurred from zero to one day after case presentation. The zone of elevated risk within one day of a primary case extended to at least 340m in N’Djamena (like the estimate for the first five days), but only 90m in Kalemie. During this time-frame, those within 20 meters of a primary case had a 76.1-fold (Kalemie; 95%CI 53.6-111.2) and 55.4-fold (N’Djamena; 95%CI 42.3-72.4) increased risk of presenting as a cholera case compared to those elsewhere in the cities. From 75-125m from a primary case, this risk decreased to 1.9 (95%CI 0.7-3.5) in Kalemie and 5.9 (95%CI 3.8-8.7) in N’Djamena.

After excluding cases occurring the same day as a primary case (day 0), the zone of increased risk was approximately the same size as estimated when those cases were included (210 m Kalemie and 310 in N’djamena, Figures 1C/1D). During this time-period, those within 20 meters of a primary case had a 2.1-fold (Kalemie; 95%CI 1.2-3.2) and 3.9-fold (N’Djamena; 95%CI 2.7-5.2) increased risk of presenting as a case compared to those elsewhere in the cities. From 75-125m from a primary case, the risk was 1.9 (95%CI 1.1-3.0) in Kalemie and 2.0 (95%CI 1.2-3.0) in N’Djamena.

### Diminishing risk over time

In secondary analyses, we explored how the elevated risk changed with time at key distances away from primary cases’ households to better illustrate the dynamic increased risk zone. We find that at 20m, a scale likely representative of a household and/or first-degree-neighbors, significant elevated risk disappeared by 3 (Kalemie) and 6 (N’Djamena) days after the presentation of the primary case (Figure S1). At 50m from the primary case household, risk remains elevated for slightly longer (7 days). Farther away from the primary case household (e.g, 150m), we that the elevated risk period may not start until 2-3 days after case presentation although it still ends by day 5-6 (Figure S1).

## Discussion

These results reveal remarkably similar spatiotemporal patterns of cholera cases across epidemics in two African cities. We find clear evidence for zones of increased risk extending at least 200m from the household of a severe cholera case within the first five days after he/she presents for care, with most elevated risk within the first days after the case and within 100m of the household. While not as dramatic, these zones of risk persist even after excluding those cases that appear on the same day. These suggest that case-centered interventions focused within 200m around cases’ households implemented within one week of case presentation may be an efficient cholera control strategy.

While the similarity in the spatiotemporal structure between the two independent settings provides reassurance that these results reflect some shared biological or structural properties of cholera transmission, there are several limitations to these analyses. First, we relied on suspected cases who sought care at health facilities. Analyses of epidemics in similar settings have shown that the true proportion of confirmed cholera cases among suspected cases can vary widely and we expect that this would tend to dilute the risk ratios, biasing results towards the null. On the other hand, people living near suspected cases might be more likely to seek care due to concerns about cholera, which could create an upward bias in our estimates. In N’Djamena, we only collected a random subset of cases household locations at the end of the epidemic. If cases within households (or neighbors) tended to seek care close together in space and time relative to the case sampling fraction, we may have captured too few pairs of cases living near one another, thus biasing our estimates of spatial risk towards unity. While there were likely people who did not seek care with mild and asymptomatic cholera, these people tend to be less infectious due to the lower bacteria concentration in their stool and the lower volume of stool produced. If these people were randomly distributed with respect to their spatiotemporal distance from another case, our estimates would vary little. However, if there was spatiotemporal clustering in mildly-symptomatic or asymptomatic cases, our results could be biased in either direction. Clustering of disease risk factors at different scales than that of transmission may influence our estimates, however the similarity between Kalemie and N’Djamena provides some reassurance that these results are generalizable to similar settings. Finally, given that we are dividing our data into spatial and temporal windows, the sample size can get small and we may not have the power to detect low levels of elevated risk. The true zones of increased risk may be larger than suggested by the 95% confidence intervals of the tau-statistic.

Case-area targeted interventions are not a new concept and have been implemented for diseases like polio and smallpox. In many cholera epidemics, case-area targeted interventions, ranging from hygiene promotion to antibiotic prophylaxis, are part of the standard protocol, although they are rarely documented or evaluated in published literature.^13, 14^ Careful reevaluation of the timing, extent and type of case-area targeted interventions are warranted. These interventions may not be ideal across all settings and identifying when (e.g, lul-periods^15^) and where they may have the biggest impact are key.

These results shed new light on the small-scale spatial structure of cholera transmission and point towards the possibility of conducting effective and efficient targeted interventions in an urban cholera epidemics. While the effectiveness of case-area targeted interventions will depend both on the types of interventions and the speed at which they are delivered, this work serves as rational for their use.

---

**Figure S1.**
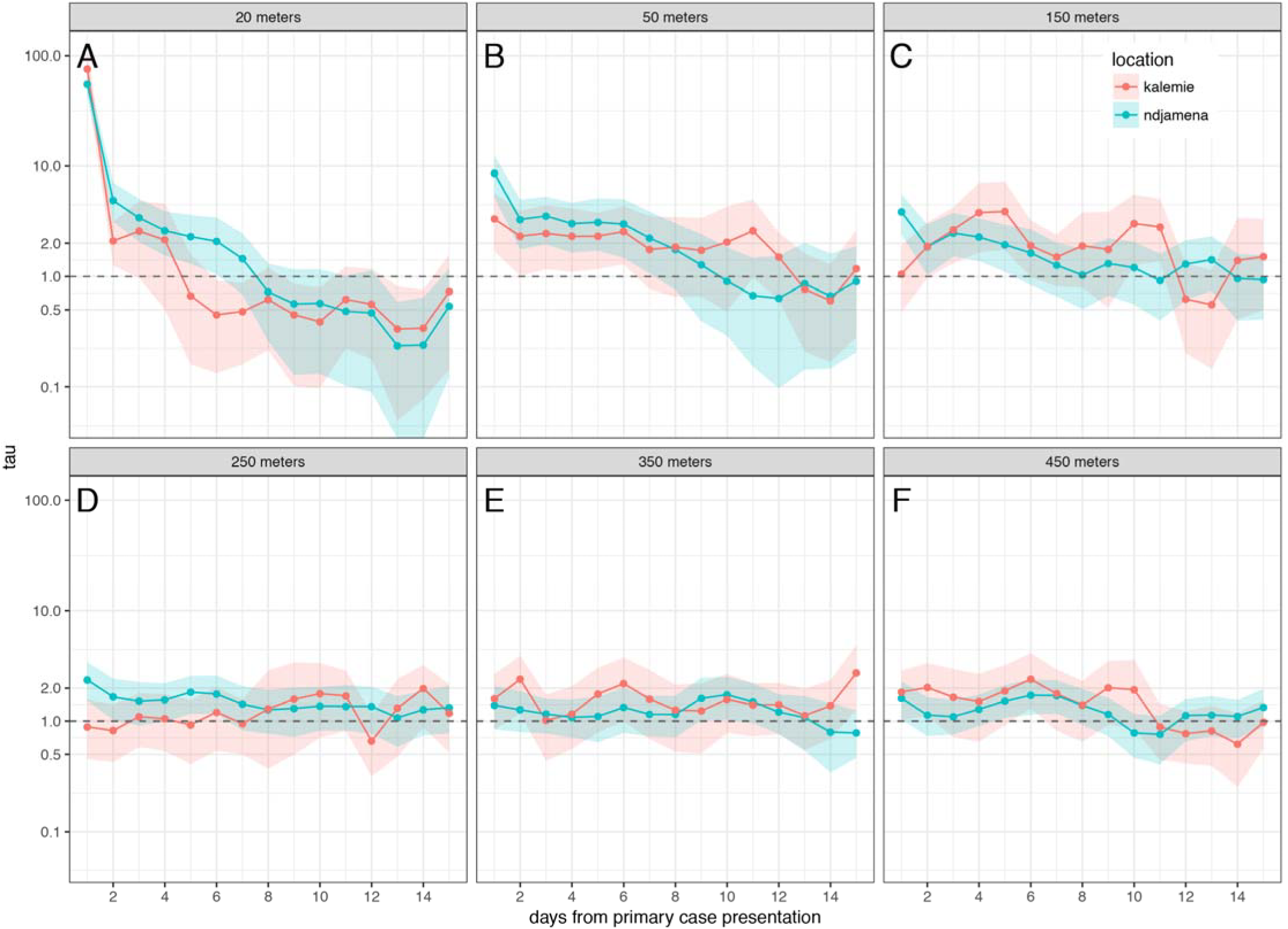
Estimates of the relative risk of the next cholera case occurring at different distances from a primary case (panels illustrate 20-200 meters) compared to the risk of the case occurring anywhere in the population by time from primary case presentation (y-axis). N’Djamena estimates and 95%CIs are shown in green and those from Kalemie are shown in orange.

**Table S1.** Median extent of the zone of increased risk based on the (bootstrap) distribution of maximum distances where \tau was greater than 1.2. 95% confidence intervals represent the 2.5^th^ and 97.5^th^ percentiles of the bootstrap distribution.

|  | Median extents of<br>zone of increased<br>risk (days 0-5),<br>95% CI | Median extents of<br>zone of increased<br>risk (days 1-5),<br>95% CI |
| --- | --- | --- |
| Kalemie | 240m (80-480) | 250m (80-590) |
| N'Djamena | 365m (290-810) | 315m (155-430) |

## References

1. Abubakar A, Azman AS, Rumunu J, et al. The First Use of the Global Oral Cholera Vaccine Emergency Stockpile: Lessons from South Sudan. 2015;12(11):e1001901.

2. Bagcchi S. Cholera in Iraq strains the fragile state. The Lancet Infectious Diseases 2016;16(1):24–5.

3. Abubakar A, Almiron M, Charito A, et al. Cholera, 2014. Weekly Epidemiological Record 2015;90(40):517–44.

4. Azman AS, Luquero FJ, Rodrigues A, et al. Urban cholera transmission hotspots and their implications for reactive vaccination: evidence from Bissau city, Guinea bissau. PLoS Negl Trop Dis 2012;6(11):e1901.

5. Rebaudet S, Sudre B, Faucher B, Piarroux R. Cholera in coastal Africa: a systematic review of its heterogeneous environmental determinants. J Infect Dis 2013;208 Suppl 1:S98–106.

6. Parker LA, Rumunu J, Jamet C, et al. Neighborhood-targeted and case-triggered use of a single dose of oral cholera vaccine in an urban setting: Feasibility and vaccine coverage. PLoS Negl Trop Dis 2017;11(6):e0005652.

7. Santa-Olalla P, Gayer M, Magloire R, et al. Implementation of an alert and response system in Haiti during the early stage of the response to the cholera epidemic. Am J Trop Med Hyg 2013;89(4):688–97.

8. Morris JG. Cholera-modern pandemic disease of ancient lineage. Emerg Infect Dis 2011;17(11):2099–104.

9. Sugimoto JD, Koepke AA, Kenah EE, et al. Household Transmission of Vibrio cholerae in Bangladesh. PLoS Negl Trop Dis 2014;8(11):e3314.

10. Bi Q, Azman AS, Satter SM, et al. Micro-scale Spatial Clustering of Cholera Risk Factors in Urban Bangladesh. PLoS Negl Trop Dis 2016;10(2):e0004400.

11. Lessler J, Salje H, Grabowski MK, Cummings DAT. Measuring Spatial Dependence for Infectious Disease Epidemiology. 2016;11(5):e0155249.

12. Salje H, Lessler J, Endy TP, et al. Revealing the microscale spatial signature of dengue transmission and immunity in an urban population. Proceedings of the National Academy of Sciences 2012;109(24):9535–8.

13. Guévart É, Noeske J, Sollé J, Mouangue A, Bikoti J-M. Antibioprophylaxie ciblée à large échelle au cours de l’épidémie de choléra de Douala en 2004. Cahiers d'études et de recherches francophones / Santé 2007;17(2):63–8.

14. George CM, Monira S, Sack DA, et al. Randomized Controlled Trial of Hospital-Based Hygiene and Water Treatment Intervention (CHoBI7) to Reduce Cholera. Emerg Infect Dis 2016;22(2):233–41.

15. Rebaudet S, Gazin P, Barrais R, et al. The Dry Season in Haiti: a Window of Opportunity to Eliminate Cholera. PLoS Curr 2013;

